# Cardiovascular Disease-Associated Non-Coding Variants Disrupt GATA4-DNA Binding and Regulatory Functions

**DOI:** 10.1101/2024.09.19.613959

**Authors:** Edwin G. Peña-Martínez, Jean L. Messon-Bird, Jessica M. Rodríguez-Ríos, Rosalba Velázquez-Roig, Diego A. Pomales-Matos, Alejandro Rivera-Madera, Leandro Sanabria-Alberto, Adriana C. Barreiro-Rosario, Jeancarlos Rivera-Del Valle, Nicole E. Muñoz-Páez, Esther A. Peterson-Peguero, José A. Rodríguez-Martínez

## Abstract

Genome-wide association studies have mapped over 90% of cardiovascular disease (CVD)-associated variants within the non-coding genome. Non-coding variants in regulatory regions of the genome, such as promoters, enhancers, silencers, and insulators, can alter the function of tissue-specific transcription factors (TFs) proteins and their gene regulatory function. In this work, we used a computational approach to identify and test CVD-associated single nucleotide polymorphisms (SNPs) that alter the DNA binding of the human cardiac transcription factor GATA4. Using a gapped k-mer support vector machine (GKM-SVM) model, we scored CVD-associated SNPs localized in gene regulatory elements in expression quantitative trait loci (eQTL) detected in cardiac tissue to identify variants altering GATA4-DNA binding. We prioritized four variants that resulted in a total loss of GATA4 binding (rs1506537 and rs56992000) or the creation of new GATA4 binding sites (rs2941506 and rs2301249). The identified variants also resulted in significant changes in transcriptional activity proportional to the altered DNA-binding affinities. In summary, we present a comprehensive analysis comprised of in silico, in vitro, and cellular evaluation of CVD-associated SNPs predicted to alter GATA4 function.

**Graphical Abstract:** 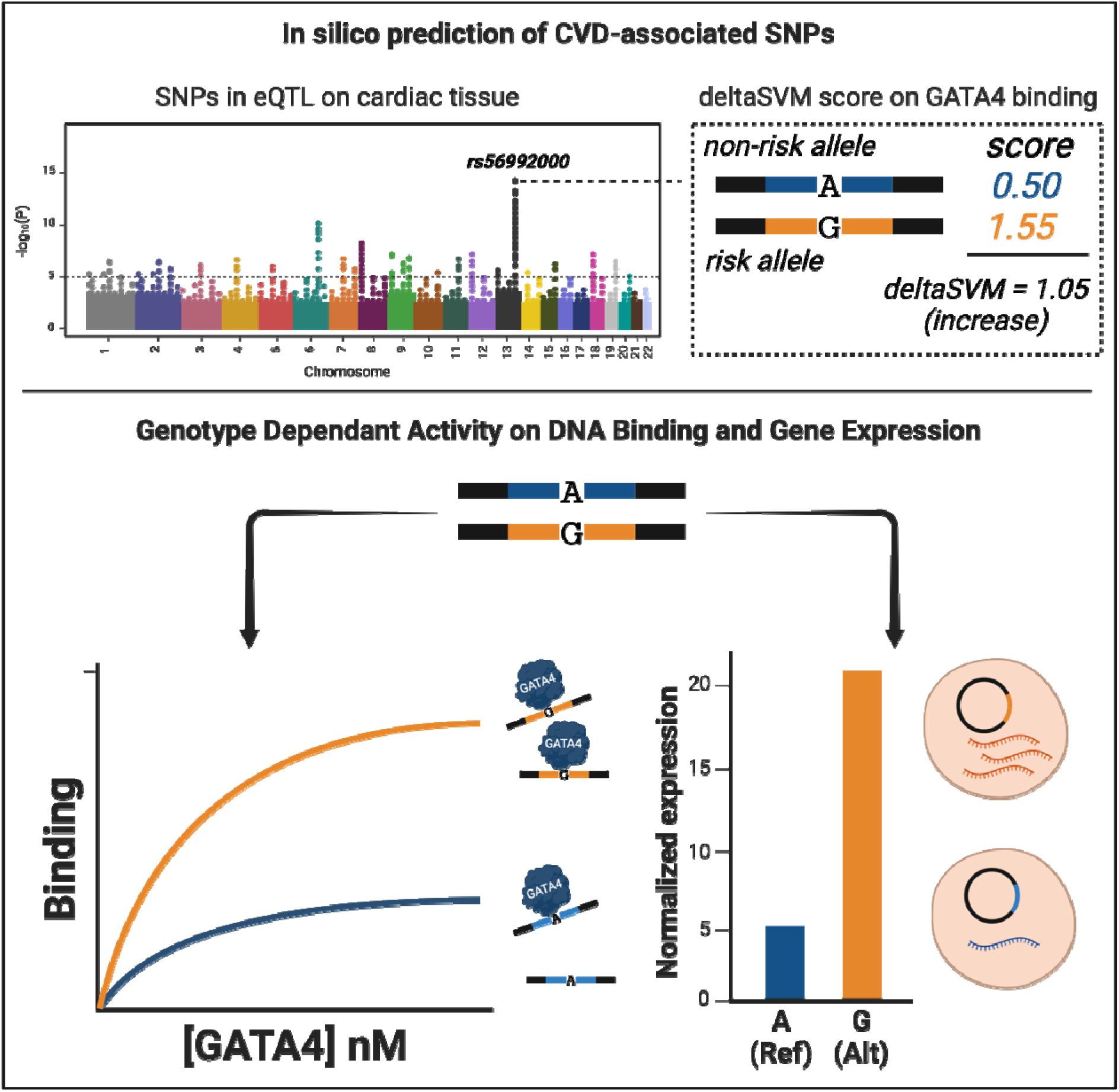

**Highlights:**

- An integrative computational approach combining functional genomics data and machine learning was implemented to prioritize potential causal genetic variants associated with cardiovascular disease (CVD).
- We prioritized and validated CVD-associated SNPs that created or destroyed genomic binding sites of the cardiac transcription factor GATA4.
- Changes in GATA4-DNA binding resulted in significant changes in GATA4-dependent transcriptional activity in human cells.
- Our results contribute to the mechanistic understanding of cardiovascular disease-associated non-coding variants impacting GATA4 function.

## Introduction

Genome-wide association studies (GWAS) have mapped the vast majority (>95%) of disease/trait-associated genetic variants within the non-coding genome. [1–6] Advancements in DNA sequencing technologies have enabled the identification of thousands of genetic variants associated with many of the leading causes of death, including diabetes, cancer, and cardiovascular diseases (CVDs). [7–15] Disease-associated variants within the non-coding regions of the genome, such as *cis*-regulatory elements (CREs, e.g., promoters and enhancers), can alter protein-DNA interactions and gene expression. [16–20] For instance, previous work has revealed that non-coding variants within CREs alter transcription factor (TF)-DNA binding [21–24], which also disrupt promoter-enhancer interactions [25,26], recruitment of co-factors [27], and regulation of gene expression [28–30]. Thus, researching how non-coding variants alter TF-DNA interactions is crucial to understanding the etiology of multiple human diseases.

GATA-binding protein 4 (GATA4) is a cardiac TF from the GATA-type zinc finger family (InterPro IPR000679) required for heart development in mammals. [31–35] During early embryonic stages, *GATA4* is expressed to initiate cardiogenesis and promote the formation of crucial structures, such as the heart tube, atrium, and ventricles. [35,36] GATA4 has a consensus sequence of 5’-`AGATAA` −3’ and interacts with other TFs (e.g., TBX5, NKX2-5, FOG2) to regulate the expression of cardiac structure genes, like the *atrial natriuretic factor (ANF)*. [33,37–39] Haploinsufficiency of *GATA4* has been strongly linked to multiple types of congenital heart diseases (CHDs), such as atrial/ventricular septal defect (ASD/VSD) and tetralogy of Fallot (TOF). Coding and non-coding variants that alter GATA4 and other members of the GATA TF family have been implicated in CHDs. [40–44] For instance, variants near the promoter of *TBX1* were previously described to impair gene activation by GATA6, implicating a functional role in conotruncal heart disease patients. [45] However, with the overwhelming and increasing number of GWAS variants reported to date (>500,000), prioritizing and validating candidate variants remains challenging.

In this work, we implemented a machine learning model to prioritize non-coding single nucleotide polymorphisms (SNPs) predicted to alter GATA4-DNA binding. Non-coding SNPs were prioritized based on **i)** association with a cardiovascular disease or cardiovascular traits in the GWAS catalog [46], **ii)** variants that occur in DNase I genomic footprints (DGF) within putative cardiac enhancers and [47,48], **iii)** genotype-dependent expression patterns in cardiac tissue (data from the GTEx portal [49]), and **iv)** support vector machine (SVM)-based model predictions to alter GATA4-DNA binding. We trained a large-scale gapped k-mer SVM (LS-GKM-SVM) with chromatin immunoprecipitation followed by DNA sequencing (ChIP-seq) data from GATA4 in human-induced pluripotent stem cells-derived cardiac progenitors (hiPSC-CPs). CVD-associated SNPs were then scored to predict changes in GATA4-DNA binding between the CVD risk-and the non-risk allele. Four variants in expression quantitative trait loci (eQTL) in cardiac tissue were selected for in vitro validation: two variants predicted to increase (rs1506537 and 56992000) GATA4-DNA binding affinity, and two variants predicted to decrease (rs2941506 and rs2301249) GATA4-DNA binding affinity. The functional impact of the four CVD-associated SNPs was validated through electrophoretic mobility shift assay (EMSA) and dual-luciferase reporter gene assay. All four variants resulted in changes in DNA-binding affinity and gene expression. In short, our combined computational and experimental approach can be applied to identify and validate non-coding variants associated with human diseases.

## Methods

### Data

ChIP-seq peaks for GATA4 cardiac progenitor cells (SRX9284038) were downloaded from ChIP-Atlas. [50] DNase I hypersensitivity footprints for fetal heart tissue (left atrium, right ventricle), heart fibroblast, and differentiated cardiomyocytes were obtained from ENCODE (ENCSR764UYH) [48]. Putative heart enhancers were downloaded from the supplementary files from Dickel et al. [47] Disease or trait-associated SNPs were downloaded from the GWAS catalog (gwas_catalog_v1.0-associations_e0_r2023–11–29.tsv). [46]

### LS-GKM SVM Model Training and Scoring

LS-GKM SVM model was implemented to predict changes in TF-DNA binding affinity for GATA4 on CVD-associated SNPs. [51,52] The LS-GKM model was trained using the top 1,000 ChIP-seq peaks (SRX9284038) with the highest intensity using the *gkmtrain* function. The following parameters were used to train the model using a 5-fold cross-validation: word length (l) = 11 and the number of informative positions (k) = 7 (gkmtrain -x 5 -L 11 -k 7 -d 3 -C 1-t 2 -e 0.005). Model performance was assessed via area under the receiver operator characteristic (AUROC) and precision-recall (AUPRC) curves using the gkmSVM package in R. The *gkmpredict* function was used to score reference (ref) and alternate (alt) allele 19-mer genomic sequences centered at the SNP. *deltaSVM* scores were calculated by subtracting the ref allele score from the alt allele score. [53] Position Weight Matrices (PWM) and DNA logos were generated by scoring every possible 11-mer and feeding the top 1,000 highest scoring 11-mer sequences into Multiple Em for Motif Elicitation (MEME) [54] web-based tool using default parameters to generate a logo.

### PWMScan Scoring

Changes in TF-DNA binding affinity for GATA4 resulting from CVD-associated SNPs were also predicted using PWMScan [55], a web-based tool for scoring genomic sequences using a PWM Model. Human GATA4 PWM MA0482.3 from the JASPAR [56,57] database was downloaded and inputted into the PWMScore Custom Weight Matrix interface. The input data sets scored by PWMScore consisted of FASTA files containing either reference or alternate 40 bp sequences centered on the CVD-associated SNPs. The predicted impact was determined by calculating the difference between ref and alt sequence PWM scores.

### Identification of CVD-associated SNPs and Selection for in vitro Validation

Variants associated with cardiovascular disease or trait were filtered using the “DISEASE/TRAIT” column with the *grepl* function as previously described [58]. To prioritize SNPs in genomic elements relevant to cardiac biology, filtered GWAS variants were intersected with heart DNase I hypersensitive genomic footprints (DGFs) [48] that occur in putative cardiac enhancers [47]. CVD-associated variants that occur in heart DGFs in cardiac enhancers were expanded to include variants in linkage disequilibrium (LD) using the LDLinkR package. [59] LD expansion was performed in five different populations (EUR, AFR, SAS, EAS, and AMR; R^2^ > 0.8). Insertions and deletions were excluded from further analysis. CVD-associated SNPs were prioritized based on eQTL in cardiac tissue (heart atrial appendage and heart left ventricle) using data from the Genotype-Tissue Expression (GTEx) portal. [49]

### GATA4 Expression and Purification

Human GATA4 DNA-binding domain (DBD), also known as a zinc finger domain (ZF), gene (Uniprot: P43694, Met207 to Ala333) was cloned into the pET-28a(+) vector with an N-terminal 6x-Histag (Twist Bioscience) and transformed into BL21 DE3 *E. coli* strain (Millipore Sigma, 70956-4). Bacteria were cultured in 50 mL of Luria Broth (Sigma Aldrich, L3022) at 37°C for 16 hours. Afterward, a 10 mL sample of the initial culture was transferred to 500 mL of Terrific Broth (Sigma Aldrich, T0918) in a 2000 mL Erlenmeyer Flask and grown at 37 °C while shaking at 130 rpm. Once the optical density at 600 nm reached 0.5-0.8, the culture was induced with 0.4 mM of Isopropyl β-d-1-thiogalactopyranoside (IPTG; Sigma Aldrich, I6758) for 20 hours at 20 °C while shaking at 130 rpm. The bacterial culture was centrifuged (2800 xg for 5 minutes at 4°C), the supernatant was discarded, and the pellet was stored at −80 °C overnight.

Purification of the GATA4 DBD was done via Ni-NTA affinity chromatography (Qiagen, 30210). Pellets were resuspended in a 40 mL of column buffer solution (20 mM Tris-HCl pH 8.0, 500 mM NaCl, 0.2% Tween-20, 30 mM imidazole, and EDTA-free protease inhibitor (Thermo Fisher, PIA32963). Afterward, 4 mL of 5 M NaCl were added and the cells were sonicated in four 30-second cycles at 40% amplitude on ice (QSONICA, Part No. Q125). The cell lysate was centrifuged (2800 xg, 30 minutes, 4°C). The chromatography column was loaded with 2 mL of resin and washed with 1 column volume of column buffer. The supernatant was loaded into the Ni-NTA affinity chromatography column and resuspended with the resin for 1 hour at 4 °C with orbital shaking. The flowthrough was passed through the column two more times. The column was washed with 20 mL washes of column buffer with increasing imidazole concentration (30, 50, and 100 mM) were performed. The protein was eluted with 1.8 mL of elution buffer (500 NaCl, 20 mM Tris-HCl pH 8.0, 0.2% Tween-20, and 500 mM imidazole) six times, and each was stored in separate tubes. Eluted protein was transferred to 5X binding buffer (50 mM HEPES, 2.5 µM zinc acetate, 500 mM NaCl, 20% Glycerol) through buffer exchange.

GATA4 DBD purity was evaluated with SDS-PAGE using 15 well Mini-PROTEAN TGX Precast Protein Gels® (Bio-Rad). Samples were prepared with 4× loading buffer containing β-mercaptoethanol (BME) for a total volume of 20 μL (5 μL 4× loading buffer: 15 μL sample) and heated at 95 ◦C for 5 min. 15 μL were loaded onto the gel for a 1.5 h run at 100 V at room temperature. The gel was stained using ProtoStain™ Blue Colloidal Coomassie G-250 stain (National Diagnostics). For the Western Blot analysis, contents from the SDS-PAGE gel were transferred to a PVDF membrane using the Bio-Rad Turbo Transfer System protocol in a Trans-Blot® Turbo™ for 3 min at 25 V. Membranes were blocked using 5 % milk in 1× TBST buffer for 1 h in orbital shaking and incubated overnight with 1:10,000 dilution of Anti-His mouse monoclonal antibody (Novus Biologicals, AD1.1.10). The SDS-PAGE and Western Blot Analysis results were imaged using Azure Sapphire Bio-molecular Imager (Azure Biosystems, **Supplementary Figure 1A**).

### Electrophoretic Mobility Shift Assay

All oligonucleotides were purchased from Integrated DNA Technologies (IDT). GATA4-DNA binding was assessed using a 20 bp oligonucleotide centered on the SNP with a 20 bp constant sequence downstream for primer extension. The primer was modified with IRDye® 700 fluorophore at the 5’-end. Incorporation of the fluorophore into dsDNA was done through a primer extension reaction using EconoTaq polymerase (Lucigen, 30035–2) and purified using the QIAquick PCR purification kit (Qiagen 28106). Binding reactions were conducted in 1X binding buffer (10 mM HEPES, 0.5 µM zinc acetate, 100 mM NaCl, 4 % Glycerol) and 5 nM fluorescently-labeled dsDNA. Binding reactions were incubated for 30 minutes at 30 °C, followed by 30-minute incubation at room temperature. Samples were loaded into a 6% polyacrylamide gel in 0.5x TBE (89 mM Tris, 89 mM boric acid, 2 mM EDTA, pH 8.4). The gel was pre-ran at 72 V for 15 minutes, loaded at 30 V, and ran at 120 V for 1 hour at 4°C. Gels were imaged using Azure Biosystem Sapphire Biomolecular Imager using 658 nm excitation and 710 nm emission.

Binding curves were generated by quantifying the pixel intensity in each DNA band using ImageJ. [60] Background intensities obtained from blank regions of the gel were subtracted from the band intensities. The fraction of bound DNA was determined using **Equation 1**. The fraction of bound DNA was plotted versus the TF concentration. Binding curves were fitted by “one-site specific binding” non-linear regression using Prism software.

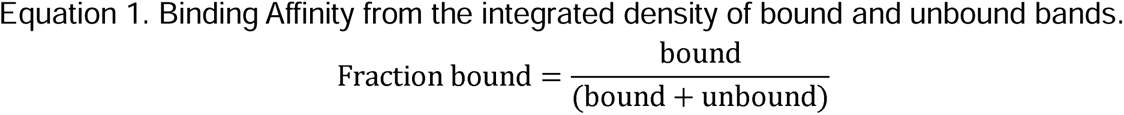

### Dual-Luciferase Reporter Gene Assay

Genomic 60 bp sequences centered on the specific SNPs were purchased from Integrated DNA Technologies (IDT) with standard desalted purification. Sequences were cloned upstream a minimal promoter into firefly luciferase vector, pGL4.23 (Promega E8411) by Gibson Assembly (NEB, E2611L). Primers and sequences used for cloning are listed in **Supplementary Table 1**. Clones were verified by whole plasmid sequencing using Oxford Nanopore (Plasmidsaurus Inc.).

HeLa Cervical Adenocarcinoma Human (HeLa) cells (ATCC–CCL-2) were grown in Eagle’s Minimum Essential Medium (EMEM) (ATCC - 30-2003) with 10% Fetal Bovine Serum (FBS) at 37 °C and 5% CO_2_. Reverse transfection was done using a ratio 3:1 volume-to-mass ratio of FuGENE6 (0.3 µL reagent: 100 ng of DNA per well, Promega E2691). The transfection mix included the following vectors: (1) pGL4.23 (Promega E8411), (2) pNL1.1.CMV [Nluc/CMV] (Promega N1091) as a transfection control and (3) GATA4 from pTwist-CMV-Beta-Glob-WPRE-Neo-3xFLAG-GATA4 (Twist Bioscience). The amount of reporter DNA (pGL4 + pNL1.1) was maintained at 82.5 ng total. 50 ng of plasmids coding for GATA4 and Green Fluorescent Protein (GFP) were transfected into HeLa cells. To confirm GATA4 expression, we performed Western Blot analysis; 600,000 cells were seeded into 6-well culture plates (Sigma-Aldrich) and transfected with 1,200 ng or 3,000 ng of GATA4 plasmid. Cell lysates were prepared with 1X RIPA Lysis Buffer (Rockland), including protease and phosphatase inhibitors, using cell scrapers (Sigma-Aldrich). The total amount of protein was calculated using the Pierce BCA Protein Assay Kit (Thermo Fisher), and 50 ug of the total lysate was loaded onto the SDS-PAGE gel. After transferring to the membrane, Anti-FLAG 1:1,000 (Sigma-Aldrich) was used to detect GATA4 expression, and Anti-GAPDH 1:1,000 (Cell Signaling) as the loading control (**Supplementary Figure 1B**).

For reporter assay, seeding was done with 20,000 cells per well in Corning® 96 Well White Polystyrene Microplates flat bottom clear, white polystyrene (TC-treated) (CCSI - CLS3610). Twenty-four hours after reverse transfection, cells were lysed according to the Nano-Glo Dual-Luciferase Reporter Assay System’s protocol (Promega N1610). First, 80 µl of ONE-Glo™ EX Reagent were added to the culture medium in each well, incubated for 3 minutes while shaking at 300 rpm. Firefly luciferase activity was measured using the Tecan plate reader Infinite 200 Pro (Tecan). Then, 80 µl of NanoDLR™ Stop & Glo® Reagent were added to each well, incubated for 10 minutes, shaking at 300 rpm and NanoLuc luciferase activity was measured. The integration time for both measurements was 0.5 second. Relative luciferase activity was obtained by dividing firefly luminescence by nanoluc luminescence (luc2/nanoluc) followed by normalization to the control, which only included the reporter plasmids. All graphs were constructed using GraphPad Prism software. Each assay was performed in triplicates with at least two biological replicates.

## Results

### Prioritization of CVD-Associated SNPs impacting GATA4 binding

To identify potential causal SNPs relevant to CVD biology, we first identified 13,982 CVD-associated variants from the GWAS catalog. CVD-associated variants were intersected with known DNase I hypersensitive genomic footprints (DGFs) from multiple heart tissues [48] within putative cardiac enhancers active during heart development and the adult heart [47], resulting in 1,535 CVD-associated variants within putative cardiac regulatory elements. These variants were expanded to include variants in linkage disequilibrium (LD) from multiple populations (EUR, AFR, SAS, EAS, and AMR; R^2^ > 0.80) that have been linked to genotype-dependent expression (eQTLs) in cardiac tissue, resulting in 14,218 unique variants (full list of variants in **Supplementary File 1**). From this final list of variants, 792 genes were identified with genotype-dependent activity in the heart atrial appendage and left ventricle. The computational pipeline to identify CVD-associated SNPs is illustrated in **Figure 1A**, and a full list of differentially expressed genes is available in **Supplementary File 1**.

**Figure 1:**
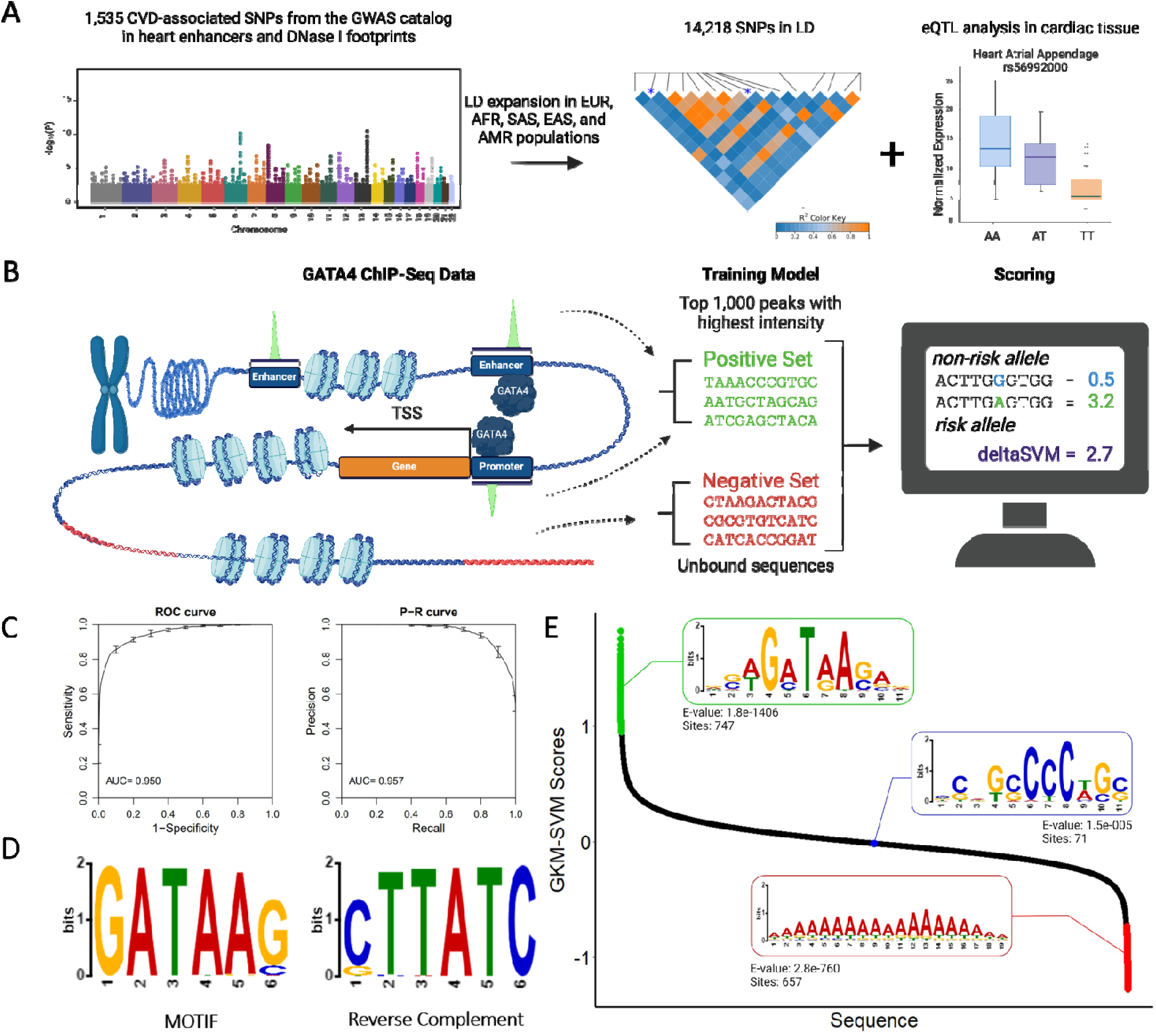
In silico approach to select CVD-associated SNPs altering GATA4-DNA binding. **A)** Pipeline to identify CVD-associated SNPs in eQTL in cardiac tissue. **B)** Schematic of model training with GATA4 ChIP-seq data from hiPSC-CPs and scoring of CVD-associated SNPs. **C)** Performance parameters of LS-GKM model. **D)** PWM logo of top 1,000 score of every possible non-redundant 11 bp oligomer. **E)** GATA4 GKM-SVM scores for ∼520,000 19-mer sequences within cardiac DGF in putative cardiac enhancers. PWM logo was generated for the top 1000 (green), middle 1000 (blue), and bottom (red) 1,000 scoring sequences.

We trained a large-scale gapped k-mer support vector machine (LS-GKM-SVM) model [51,52] to predict the effect of single nucleotide variant in the DNA-binding affinity of GATA4, a cardiac TF crucial for heart development. The LS-GKM model was trained using GATA4 ChIP-seq data from cardiac progenitor cells (SRX9284038). The 1000 top-scoring ChIP-seq peaks were used as the positive training set, while unbound sequences of the same length, GC content, and chromosome were used as the negative training set. Training of the LS-GKM model and sequence scoring is illustrated in **Figure 1B**. The best-performing LS-GKM model had satisfactory performance parameters as determined through AUROC (0.95) and AUPRC (0.96) values (**Figure 1C**). To test the model, we scored all possible 2,097,152 nonredundant 11 bp oligomers (11-mers). The 1000 top-scoring 11-mers were used to generate a PWM using MEME [54] and observed DNA-binding motifs in agreement with those previously described for GATA4 [39,61] (**Figure 1D**). Chromatin accessibility (e.g., DNase-seq and ATAC-seq) and epigenetic modifications associated with active enhancers (H3K27ac & H3K4me1) are frequently used to identify candidate active regulatory elements in specific tissues. To identify potential cardiac regulatory elements bound and regulated by GATA4, we scored 519,540 19-mer genomic sequences that reside within heart DGF in putative cardiac enhancers using the LS-GKM-SVM GATA4 model. A de novo motif discovered in the top 1,000 cardiac regulatory elements matched the known GATA4 motif (**Figure 1E**, full list of sequences and scores in **Supplementary File 1**). Whereas elements with the lowest 1000 scores did not retrieve GATA4 motifs.

### CVD-associated SNPs alter GATA4-DNA Binding

We selected four SNPs among the highest predicted impact on GATA4-DNA binding. Variants rs1506537 (largest predicted increase) and rs56992000 were predicted to increase GATA4-DNA binding, whereas rs2941506 and rs2301249 (largest predicted decrease) are predicted to decrease GATA4-DNA binding (**Figure 2A**). When evaluated for in vitro binding, we observed significant differences in GATA4-DNA binding between the reference and the alternate alleles for all four SNPs (**Figure 2B-C, Supplementary Figure 2**). As predicted by the LS-GKM SVM model, variants rs1506537 and rs56992000 showed increased GATA4-DNA binding affinity (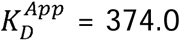 and 451.7 nM, respectively for alternate allele) compared to no binding observed for the reference allele. In both variants, the alternate allele results in a perfect match to the GATA4 cognate site (5’-`AGATAA` −3’) (Figure 2C). In agreement with LS-GKM SVM model predictions, GATA4 bound variants rs2941506 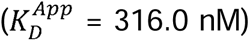 and rs2301249 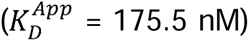, whereas almost no binding was observed for the alternate allele. Not surprisingly both variants change high-information content positions in GATA4 cognate site (**Figure 2C**).

**Figure 2:**
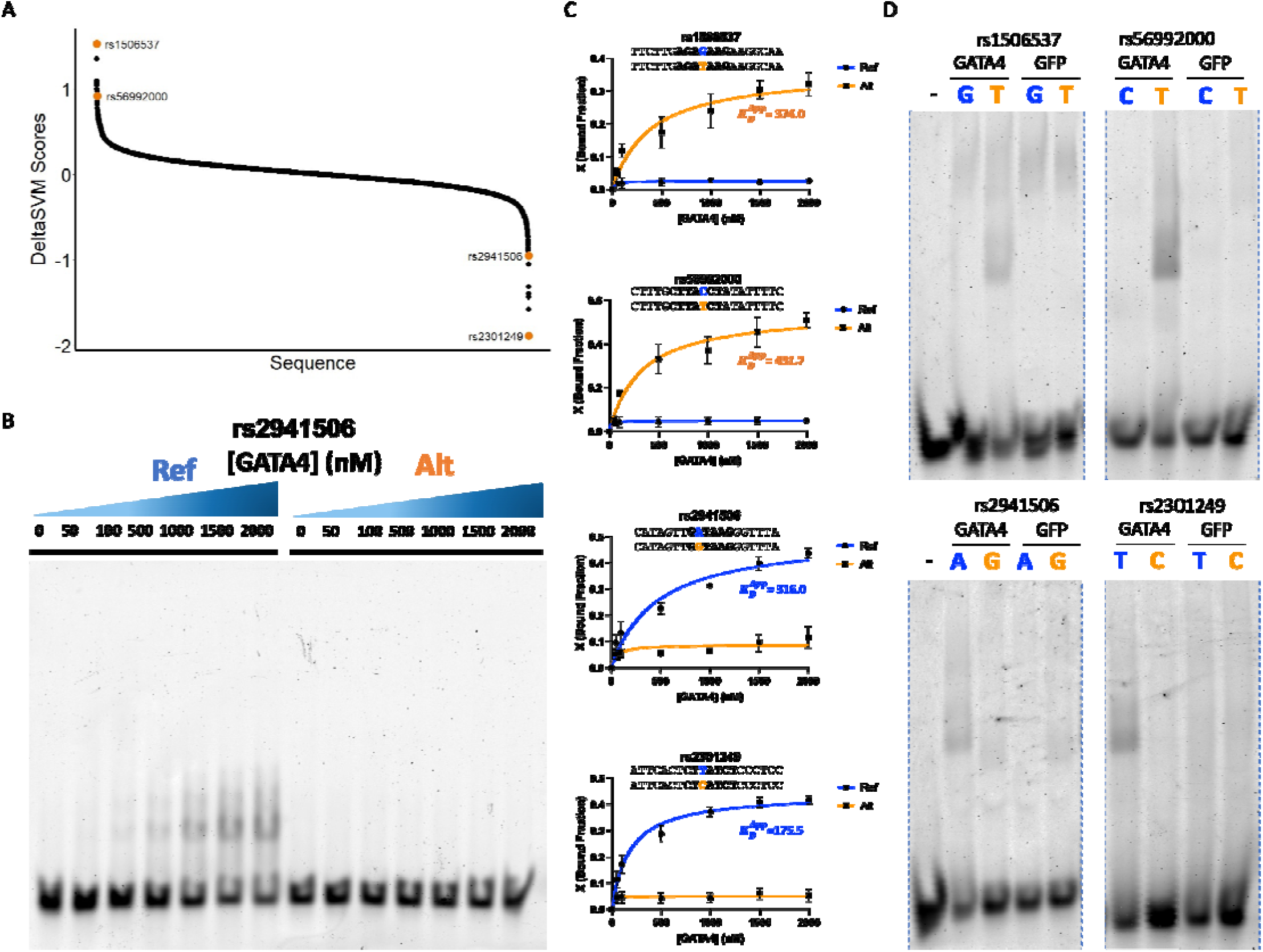
CVD-associated SNPs alter GATA4-DNA binding in vitro. **A)** GATA4 DeltaSVM scores for 9,105 CVD-associated SNPs. **B)** Representative EMSA gel for rs2941506 reference/non-effect (Ref) and alternate/effect (Alt) alleles. **C)** Binding curves of purified GATA4 DBD and CVD-associated SNPs rs1506537, rs56992000, rs2941506, and rs2301249. Experiments were performed in triplicate and binding curves show average bound fraction (X) and error bars are standard error. **D)** EMSA gels for cell lysates overexpressing full-length GATA4 and reference and alternate alleles for rs1 06537, rs56992000, rs2941506, and rs2301249. Cell lysate overexpressing green fluorescent prote n were as a negative control.

We wanted to test whether the predicted changes in binding also held for full-length GATA4 expressed in human cells. Binding with full-length proteins expressed in eukaryotic cells provide an additional layer of cellular context, such as proper folding, post-translational modifications, and binding partners such as co-factors and other TFs missing in the bacterial system. Full-length GATA4 was overexpressed in HeLa cells, and DNA binding was performed using whole-cell lysates. We observed changes in full-length GATA4-DNA binding affinity in agreement with the LS-GKM SVM model predictions (**Figure 2D, Supplementary Figure 3**). In short, binding experiments with full-length GATA4 agreed with those performed using the DNA-binding domain expressed in a bacterial system.

### Comparison of LS-GKM SVM and PWM-based Predictive Models on CVD-associated SNPs

To compare the predictions of the LS-GKM SVM model with those a Position-Weight Matrix model, we also scored the 9,105 CVD-associated SNPs using a PWM-based model. Overall, both models agreed with about half of the predictions of GATA4-DNA binding activity (whether they categorically increased or decreased binding). About 4,395 SNPs had contradictory predictions between the two models (**Figure 3A**, full list of sequence scores in **Supplementary File 1**). We selected three variants with contradictory predictions to test GATA4 binding in vitro (**Figure 3B**). Variant rs7078507 was predicted to decrease GATA4-DNA binding by the PWM-based model (deltaPWM = −901; rank 9092/9105; 0.002 percentile), but predicted to increase by the SVM-based model (deltaSVM = 0.15; rank 1687/9105; 0.81 percentile). Whereas, variants rs7372902 and rs6753736 were predicted to increase GATA4-DNA binding affinity by the PWM-based model (deltaPWM = 227; rank 81/9105, 0.90 percentile and 1197; rank 16/9105, 0.99 percentile, respectively) but predicted to decrease by the SVM-based model (deltaSVM = −0.21; rank 8209/9105 and −0.288; rank 8863/9105, respectively). Variant rs7078507 increased GATA4-DNA binding, while rs7372902 and rs6753736 decreased GATA4-DNA binding (**Figure 2C-D, Supplementary Figure 4**). For two of the three disagreeing SNPs tested, we saw notable changes in their binding affinity, a 2.4-fold decrease for rs6753736, and a 2.8-fold increase for rs7078507. In all cases, in vitro binding experiments agreed with the LS-GKM SVM-based, as determined by quantifying the bound fractions and calculating the apparent dissociation constant (K_d_) for both alleles.

**Figure 3:**
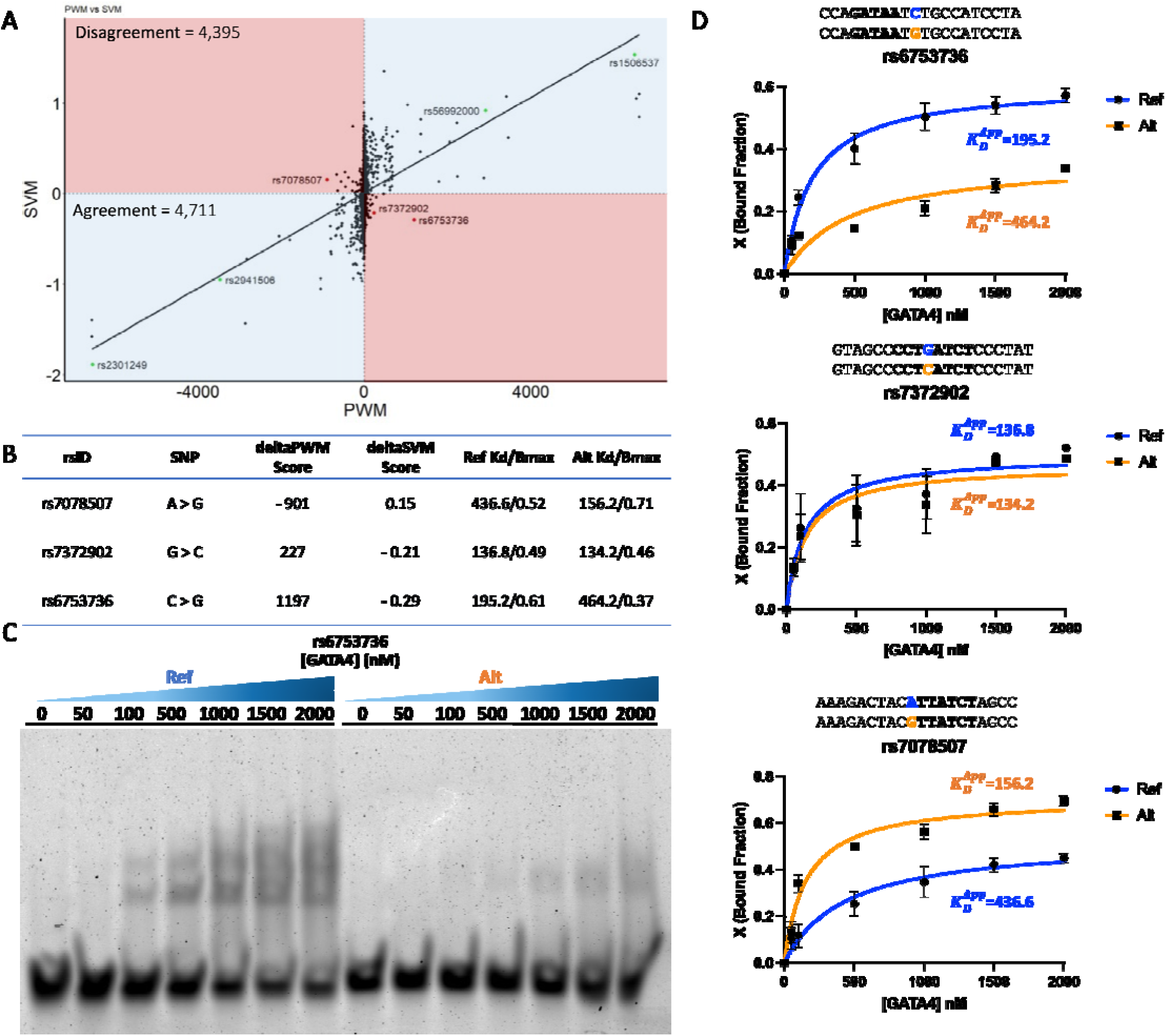
LS-GKM classifier outperforms PWM-based predictive model. **A)** Correlation of CVD-associated SNPs scored by SVM-(y-axis) and PWM-(x-axis) based predictive models. **B)** Table of example variants with contradictory binding predictions between SVM and PWM models. **C)** Representative EMSA gel for rs6753736 reference/non-effect (Ref) and alternate/effect (Alt) alleles. **D)** Binding curves of rs6753736, rs7372902, and rs7078507. Experiments were performed in triplicates and binding curves show average bound fraction (X) and error bars are standard error.

### CVD-associated SNP alter GATA4 Regulatory Activity

To determine the impact of CVD-associated SNPs on GATA4 gene regulatory activity, we performed a dual-luciferase reporter gene assay. HeLa cells were co-transfected *GATA4* plasmid along with luciferase reporter vectors with 60-bp sequences centered on variants rs2941506, rs2301249, rs1506537, and rs56992000 (**Figure 4A**). The alternate allele of variant rs2941506 resulted in a significant decrease (−1.3-fold, t-test p-value = 0.041 in luciferase activity compared to the refence allele. Whereas no significant difference in luciferase activity was observed between the reference and alternate alleles of variant rs2301249 (p-value = 0.442). Alternate alleles for variants rs1506537 and rs56992000 resulted in a higher (1.3- and 2.1-fold, t-test p-value = 0.011 and 0.0003, respectively) luciferase activity compared to their respective reference allele.

**Figure 4:**
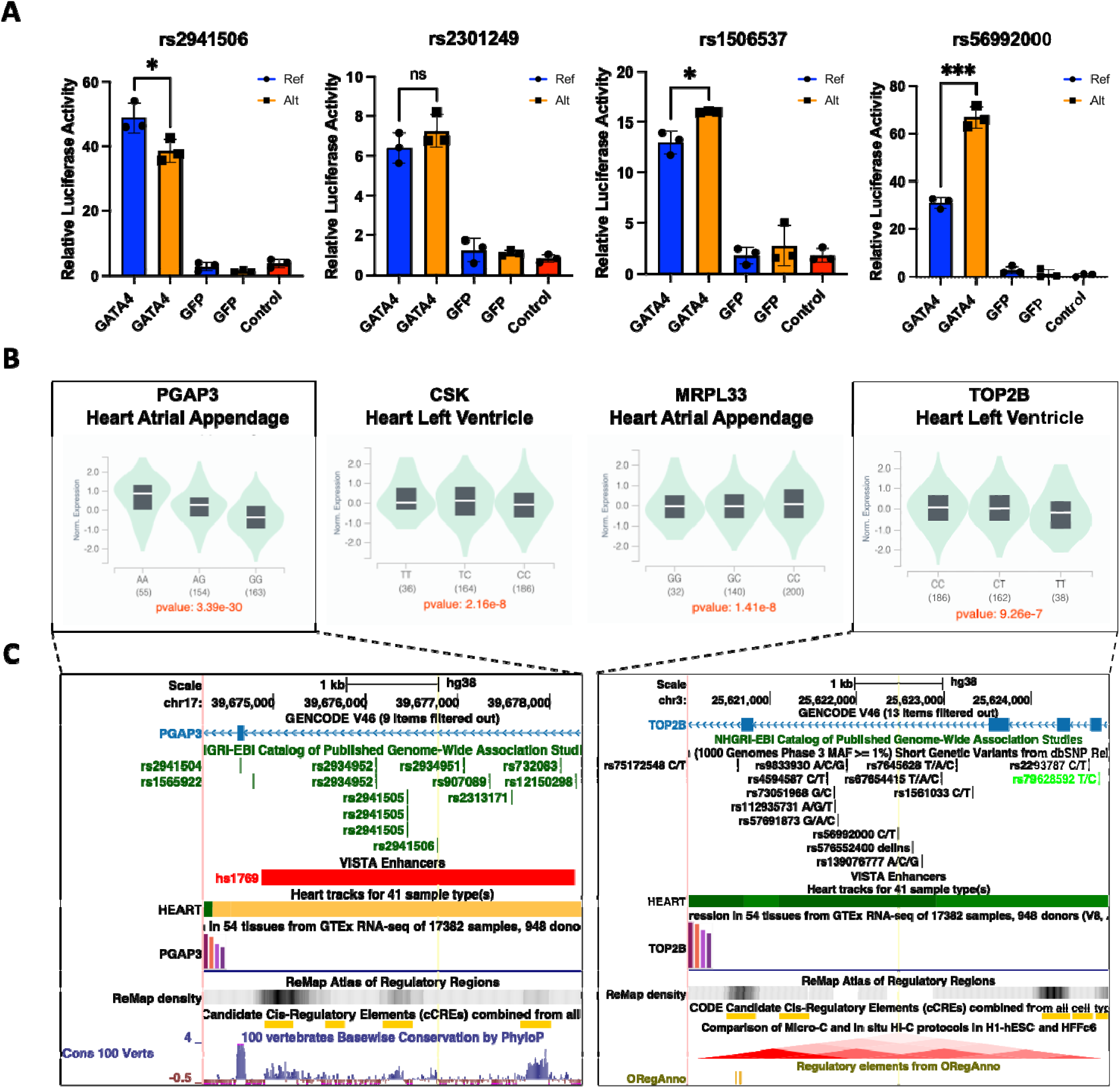
CVD-associated SNPs alter gene expression and are in eQTL in cardiac tissue. **A)** Relative luciferase activity in HeLa cells transfected with reporter plasmids containing reference (blue with black circles) and alternate (orange with black squares) of variants rs1506537, rs56992000, rs2941506, and rs2301249. **B)** Cardiac tissue eQTL analysis of *MRPL33*, *TOP2B*, *PGAP3*, and *CSK* expressed in heart atrial appendage or left ventricle when rs1506537, rs56992000, rs2941506, and rs2301249 occur, respectively. **C)** University of California Santa Cruz (UCSC) Genome Browser tracks of variants rs2941506 and rs56992000 near TAD boundaries and regulatory elements.

All four variants, rs2941506, rs2301249, rs1506537, and rs56992000, are in eQTL in cardiac tissue (**Figure 4B, Supplementary Figure 5**). Alternate alleles of variants rs1506537 and rs56992000 increased GATA4-DNA binding and reporter gene expression and are associated with increased MRPL33 expression in the heart atrial appendage and decreased *TOP2B* expression in the heart’s left ventricle, respectively. Alternate allele of variant rs2941506 decreased GATA4-DNA binding affinity compared to reference allele is associated with decreased *PGAP3* (heart atrial appendage and left ventricle), *PNMT* (heart atrial appendage), and *PPP1R1B* (heart atrial appendage), and *ERBB2* (heart atrial appendage) expression. And alternate allele of variant rs2301249, which has decreased affinity for GATA4 compared to the reference allele, was associated with decreased expression of *ULK3* in the heart atrial appendage and left ventricle and *CSK* in the left ventricle. Notably, these variants occur within regulatory elements (e.g., enhancers and promoters) within topologically associated domain (TAD) boundaries, ChIP-seq peaks of other TFs, and accessible regions in cardiac tissue (**Figure 4C, Supplementary Figures 7 and 8**), as determined by ENCODE, and near most dysregulated genes from the eQTL analysis. [62]

## Discussion

The number of disease-associated genetic variants is continuously increasing due to accessibility and advancements in DNA sequencing technologies and genome-wide association studies. It is crucial to identify causal variants for human diseases and evaluate their biochemical, cellular, and organismal implications. However, prioritizing functional/causal variants remains challenging. In this work, we employed a gapped -kmer SVM classifier to identify CVD-associated SNPs predicted to alter GATA4-DNA binding and leveraged on available functional genomic to identify SNPs with putative roles in cardiac biology. We prioritized four CVD-associated SNPs to validate their impact on biochemical interactions in vitro and gene regulation in living cells.

First, we leveraged on genomic datasets (e.g., GWAS catalog, ChIP-seq, DNase I hypersensitive genomic footprints, eQTLs, etc.) and computational tools to identify cardiovascular diseases and quantitative traits associated SNPs and variants from diverse populations (EUR, AFR, SAS, EAS, and AMR). Genetic variants were further filtered to prioritize SNPs in eQTL in cardiac tissues. We then trained a GKM SVM classifier using GATA4 ChIP-seq data from human induced pluripotent stem cell cardiac progenitors (hIPSC-CP). The GKM SVM model was assesed in silico by scoring non-redundant 11-mers and generating a PWM logo in agreement with those previously described for GATA4. Additionally, DNase I genomic footprints within cardiac enhancers were scored, identifying 747 sites potentially regulated by GATA4. Although millions of putative enhancers have been identified, the TFs that bind to them are mostly unknown. [63,64] Our approach provides a feasible strategy to identify tissue-specific regulators in silico.

We next proceeded to validate four non-coding SNPs that predicted an increase (rs1506537 and rs56992000) and decrease (rs2941506 and rs2301249) in GATA4-DNA binding affinity and are in eQTL in cardiac tissue. When evaluated through in vitro binding, variants rs1506537 and rs56992000 increased DNA-binding affinity compared to the reference sequence with almost no DNA binding even at 2,000 nM of the GATA4 DBD. Conversely, variants rs2941506 and rs2301249 resulted in a total loss of DNA binding as predicted by the LS-GKM SVM model. The in-silico predictions also held when evaluated using full-length GATA4 from human whole-cell lysates, which have additional layer of cellular context, such as proper folding, post-translational modification, and binding co-factors. Our findings suggest that the GKM SVM classifier effectively predicts changes in DNA-binding affinity for full-length and DBDs of human TFs.

To further test the GKM SVM classifier, we compared the scored CVD-associated SNPs with the predictions of a PWM-based predictive model. Overall, the GKM SVM- and PWM-based models agreed with most of the predicted changes in GATA4 binding. However, some variants had contradictory predictions between the GKM SVM and PWM models. We observed that in those cases, GKM SVM model predictions agreed with our in vitro GATA4-DNA binding measurements. Notably, two of these variants in SNPs (rs6753736 and rs7078507) are in positions flanking the GATA4 cognate site, which have been previously shown to contribute to the TF binding. [65,66] The PWM-based predictive models have several limitations, such as depending on perfect binding motifs, lacking information on complex intracellular patterns, and ignoring dinucleotide interactions. [17,67–70] In short, our data suggests that the GKM SVM model can better identify the effect in TF binding of important positions outside the TF cognate sites. [53,71,72]

Finally, we evaluated how changes in GATA4-DNA binding could alter gene expression in a cellular context through luciferase gene reporter assays. We observed significant changes in luciferase activity in agreement with changes in DNA binding for variants rs1506537, rs56992000, and rs2941506. For variant rs2301249, we observed an increase in luciferase activity even though DNA binding affinity decreased for full-length GATA4 and the ZF domain. Although not statistically significant, GATA4 has been previously reported to gain repressor functions depending on its TF binding partners and cofactors. [38,73–75] The results from luciferase reporter assays were also consistent with the expression patterns observed in eQTL analysis in cardiac tissue for the variants rs1506537 (*MRPL33*), rs56992000 (*TOP2B*), rs2941506 (*PGAP3*, *PNMT*, *PPP1R1B*, and *ERRB2*), and rs2301249 (*ULK3* and *CSK*). The *MRPL33* gene has been previously identified as a biomarker for coronary artery disease and a regulator of blood pressure. [76,77] Dysregulation of *TOP2B* and *ERRB2* has been associated with a higher risk of cardiac dysfunction and cardiotoxicity drug responses in cardiomyocytes. [78–80] Expression of *PNMT* has been implicated in several pathways involved in cardiomyocyte differentiation. [81] The long non-coding RNA PPP1R1B regulates essential genes for cardiac maturation and muscle development in mice and humans. Variant rs2941506, mapped near *PGAP3*, has been previously associated with the development of asthma and cardiovascular diseases. [82,83] In cardiovascular medicine, CSK signaling pathways are used to regulate molecules that oversee alleviating cardiac microvascular injuries, particularly in patients with diabetes. Additionally, variant rs2301249, mapped near *CSK*, has been associated with arterial and systolic blood pressure. [84] Our findings suggest that variants rs1506537, rs56992000, rs2941506, and rs2301249 are potential SNPs that can be further researched as causal variants for CVDs through GATA4 dysregulation.

In short, we implemented an LS-GKM SVM classifier to identify potential CVD-causing variants that alter GATA4-DNA interactions and regulatory functions. The SVM classifier was tested in silico to identify heart DGF within cardiac enhancers potentially regulated by GATA4. We compared two predictive models, finding that while the PWM-based model could not predict mutations outside the core binding motif, the SVM classifier successfully forecasted changes in the binding affinity of SNPs in flanking regions of TF binding sites. We further validated the predictions in vitro of four CVD-associated SNPs that drastically altered full-length GATA4 and ZF domain-DNA binding. These variants also showed changes in gene expression comparable to genes in eQTL in cardiac tissue. In summary, we present a comprehensive analysis comprised of in silico, in vitro, and cellular evaluation of CVD-associated SNPs predicted to alter GATA4 function.

## Acknowledgment

This project was supported by NIH-SC1GM127231, NSF [1736026], University of Puerto Rico Rio Piedras Institutional Funds (FIPI), Puerto Rico Science, Technology, and Research Trust, and NIH Institutional Development Award (IDeA) INBRE [P20GM103475]. EGMP, JMRR, RVR, DAPM, ACBR, LSA, and NEMP were funded by the NIH RISE Fellowship (5R25GM061151–20). JLMB and ARM were funded by NSF PR-LSAMP fellowship (HRD-2008186). DAPM was funded by NSF [IQ BIOREU 1852259]. EGPM and JMRR were funded by the NSF BioXFEL Fellowship (STC-1231306). ARM was funded by NSF REU: PR-CLIMB Program (2050493) and NIH 1T34GM145404. LSA was funded by NIH ID-GENE Fellowship (1R25HG012702–01). JMRR was funded by NSF Graduate Research Fellowship (1744619). The graphic abstract and Figure 1A-B were made using Biorender.

## Author Contribution

EGPM, JLMB, and JARM conceived and designed the study. JLMB, JMRR, RVR, ARM, LSA, ACBR, JRDV, and NEMP performed the experiments. NEMP and RVR performed recombinant GATA4 cloning, expression, and purification. JLMB, ARM, LSA, and JRDV performed EMSA. ACBR and JMRR performed dual luciferase reporter assays. EGPM and JMRR analyzed the data from the in vitro binding and gene reporter experiments. EGPM and DAPM conducted the bioinformatic analysis for LD expansion, eQTL analysis, and SVM/PWM training and scoring. EGPM and JLMB wrote the original manuscript. EGMP, ARM, JMRR, and JARM revised and edited the original manuscript. EAPP and JARM supervised the work and secured funding. All authors read and approved the manuscript.

## Competing Interests

The authors declare no competing interest.

## Supplementary Material

**Supplementary Figure 1:**
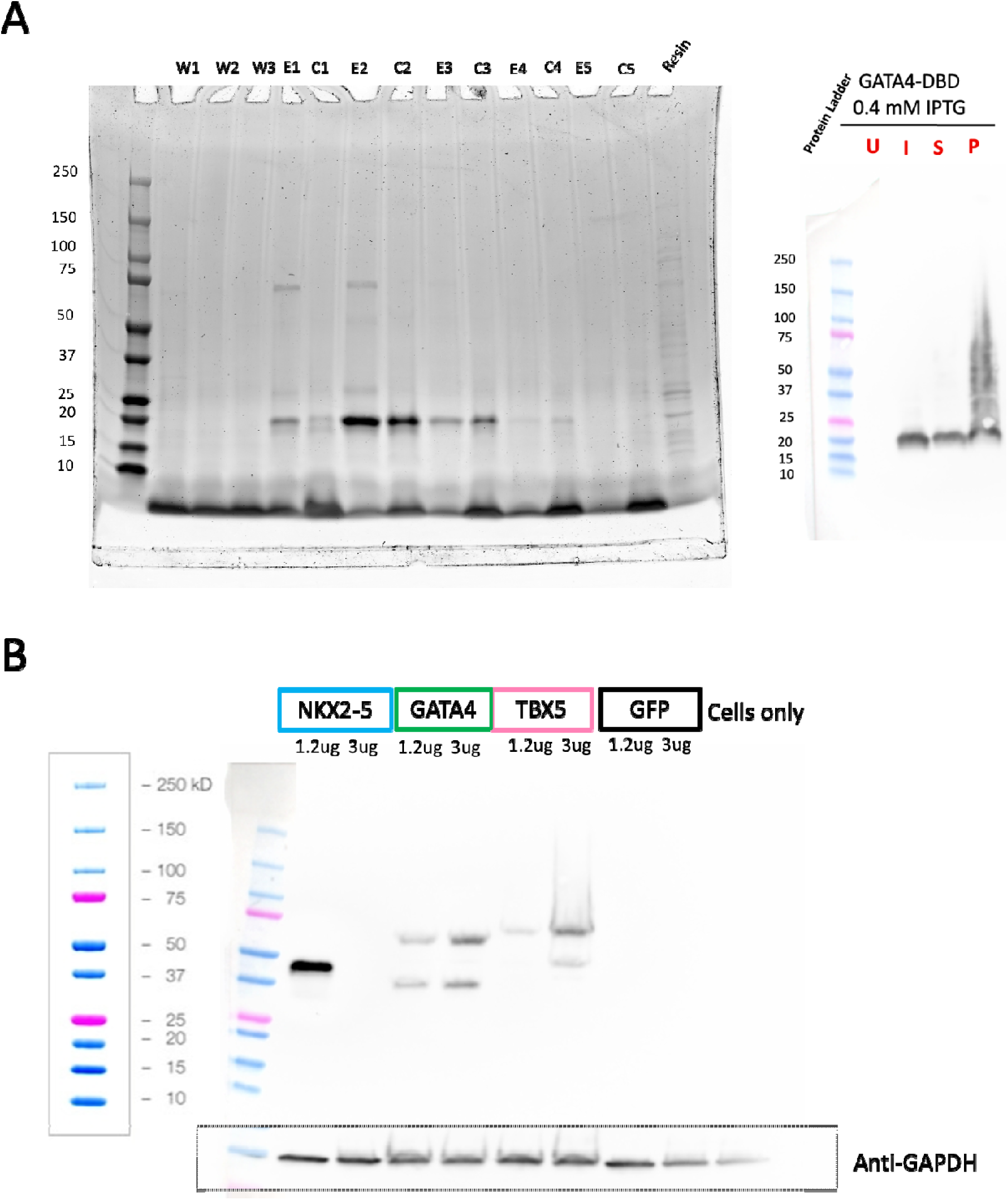
Expression and purification of GATA4 ZF. **A)** SDS-PAGE (left) and western blot (right) of GATA4 ZF (18.8 kDa). Abbreviations: wash fractions (W), eluted fractions (E), concentrated fractions (C), un-induced protein sample (U), IPTG-induced protein sample (I), supernatant fraction (S), and pellet fraction (P). **B)** Western Blot of full-length GATA4 expressed in HeLA cells (48.6 kDa).

**Supplementary Figure 2:**
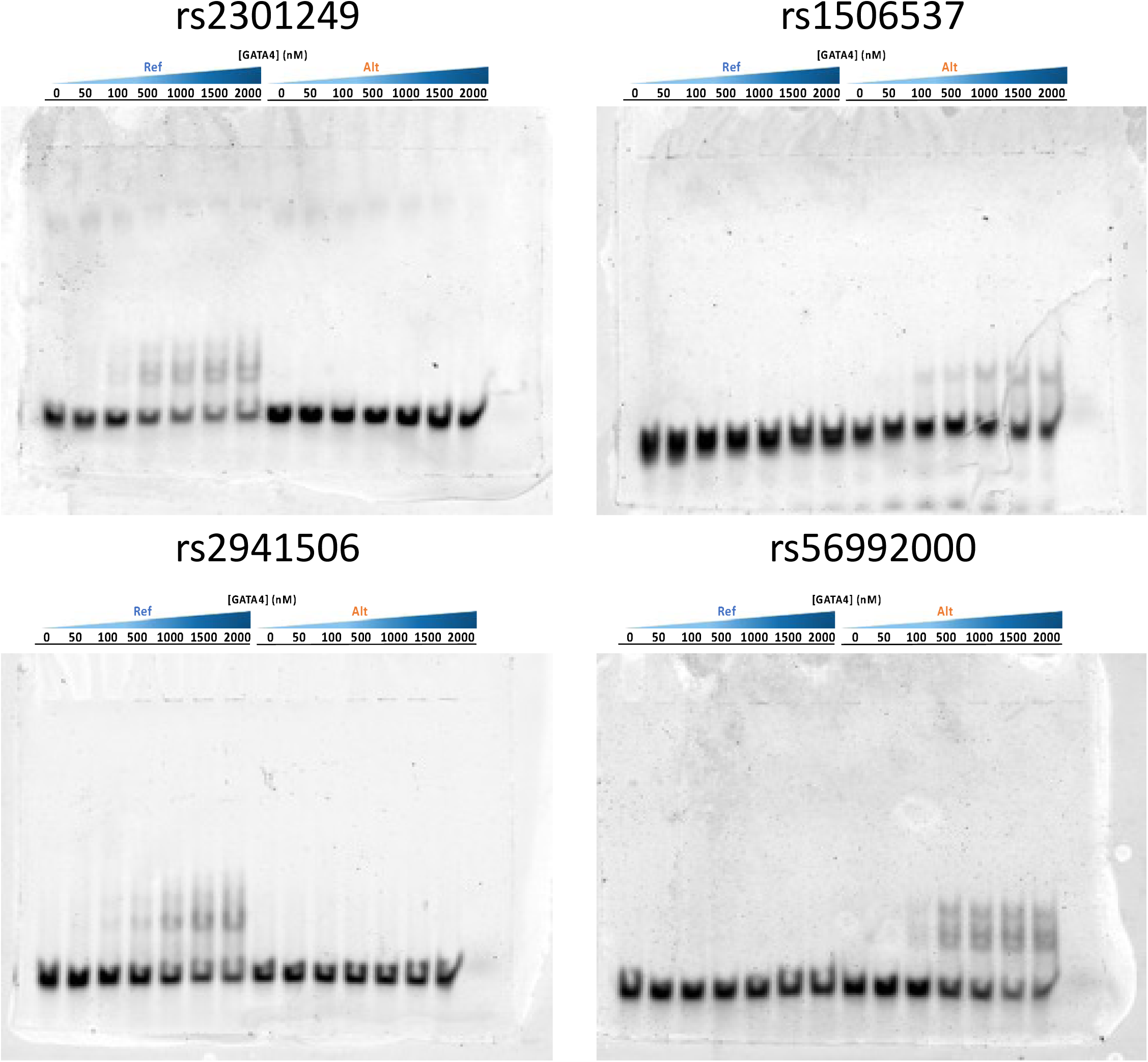
EMSA analysis of CVD-associated SNPs on GATA4 ZF domain for variants rs2301249 (top left), rs1506537 (top right), rs2941506 (bottom left), and rs56992000 (bottom right).

**Supplementary Figure 3:**
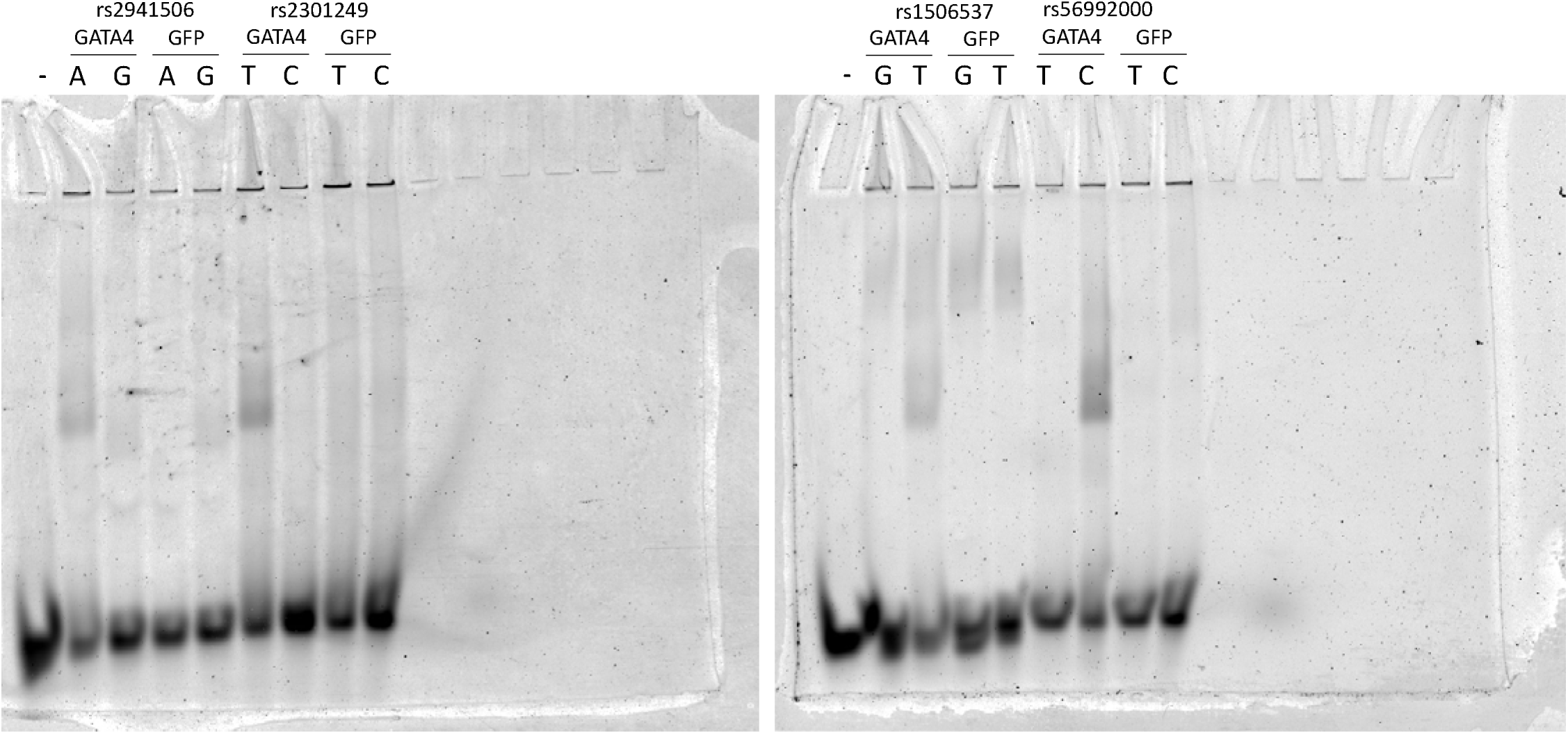
EMSA analysis of CVD-associated SNPs on full-length GATA4 overexpressed lysates for variants rs2301249, rs1506537, rs2941506, and rs56992000.

**Supplementary Figure 4:**
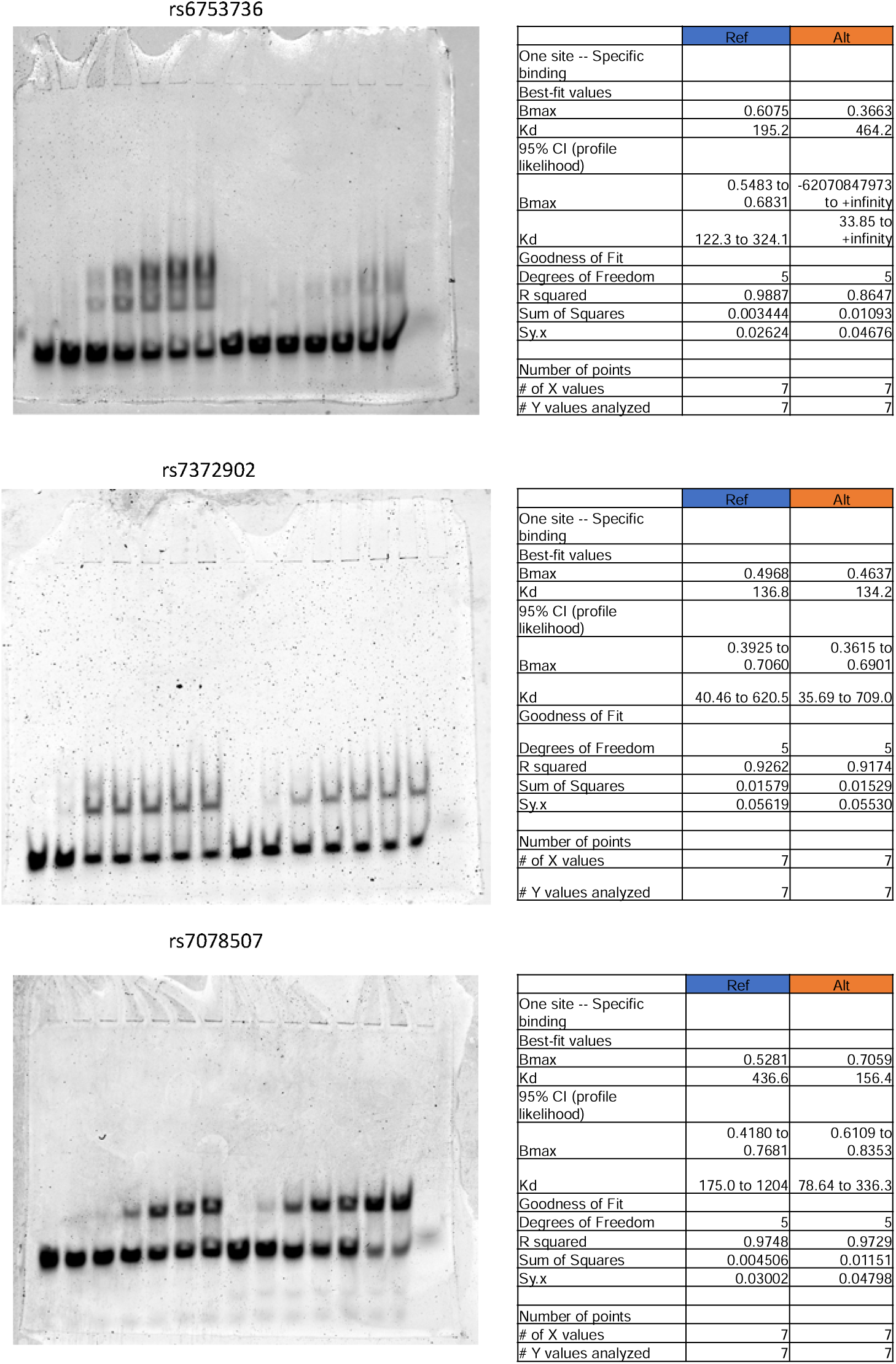
EMSA analysis of CVD-associated SNPs scored with GKM and PWM-based models. Representative EMSA for variants rs6753736 (top), rs7372902 (middle), and rs7078507 (bottom). Each EMSA was performed and analyzed in triplicates. Statistical analysis and quantification of each EMSA are on the right of each gel.

**Supplementary Figure 5:**
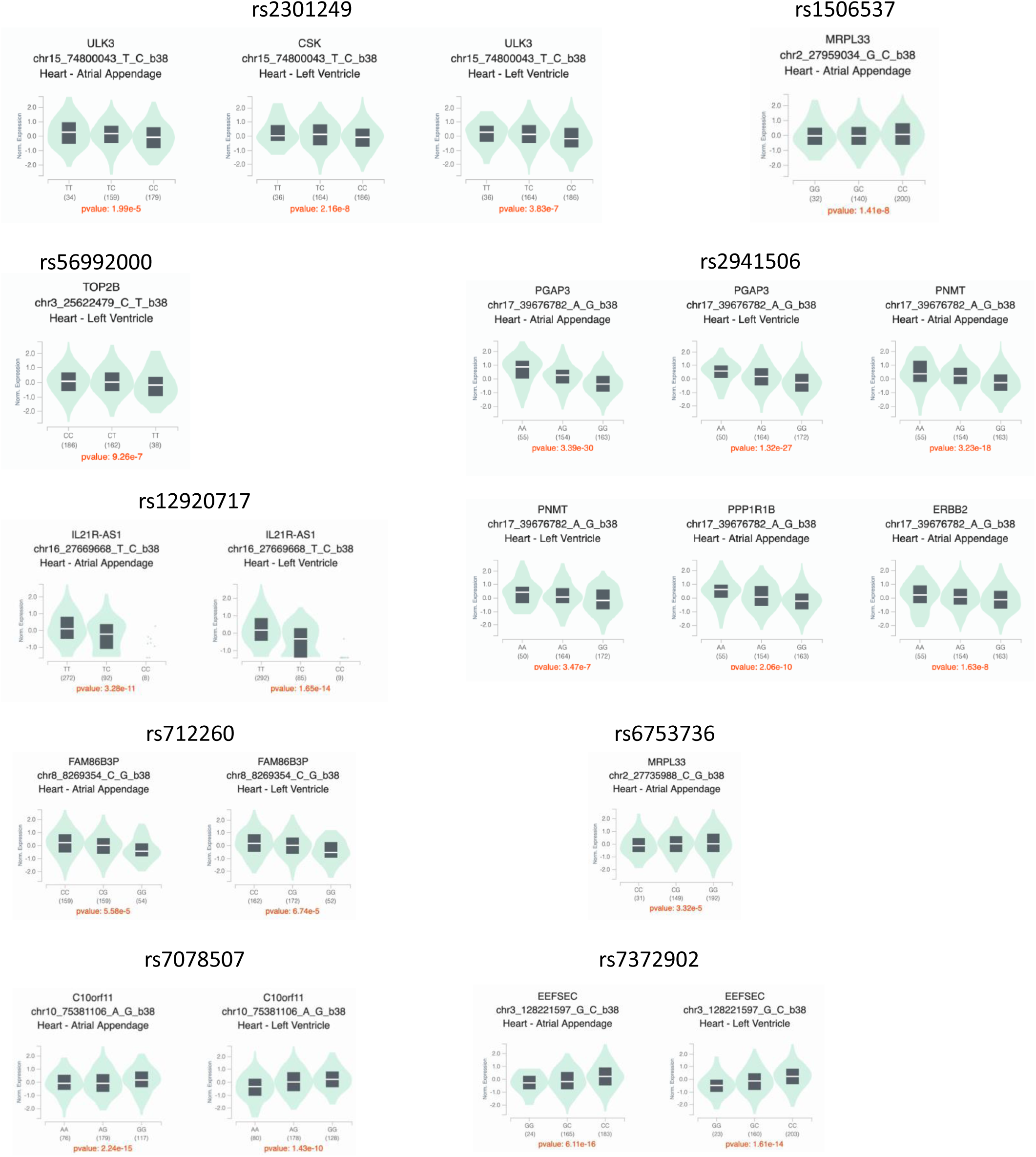
eQTL analysis of CVD-associated SNPs in cardiac tissue (heart atrial appendage and left ventricle).

**Supplementary Figure 7:**
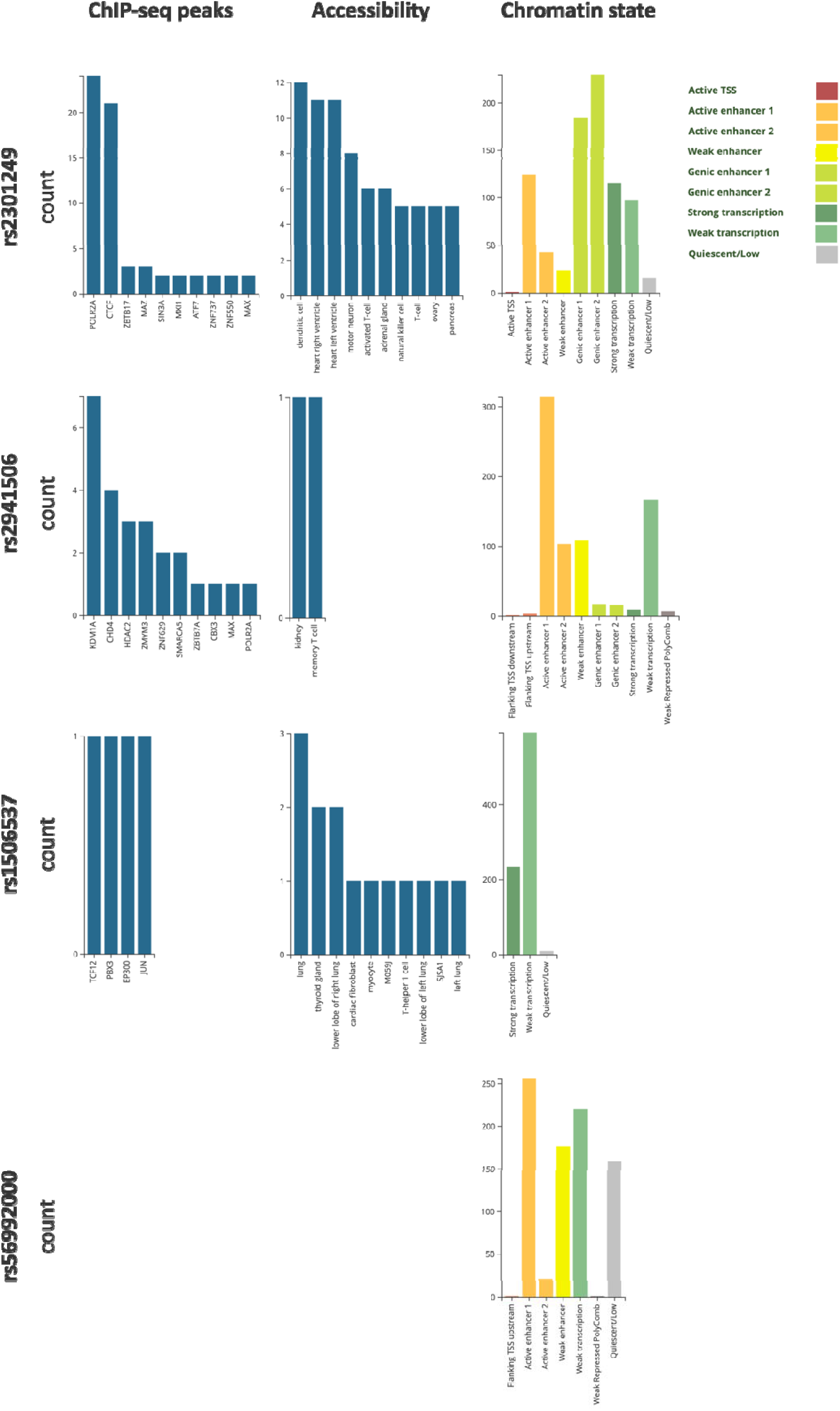
Summary of CVD-associated SNPs from Figure 2 in regulatory regions. Data was downloaded from RegulomeDB [62]

**Supplementary Figure 8:**
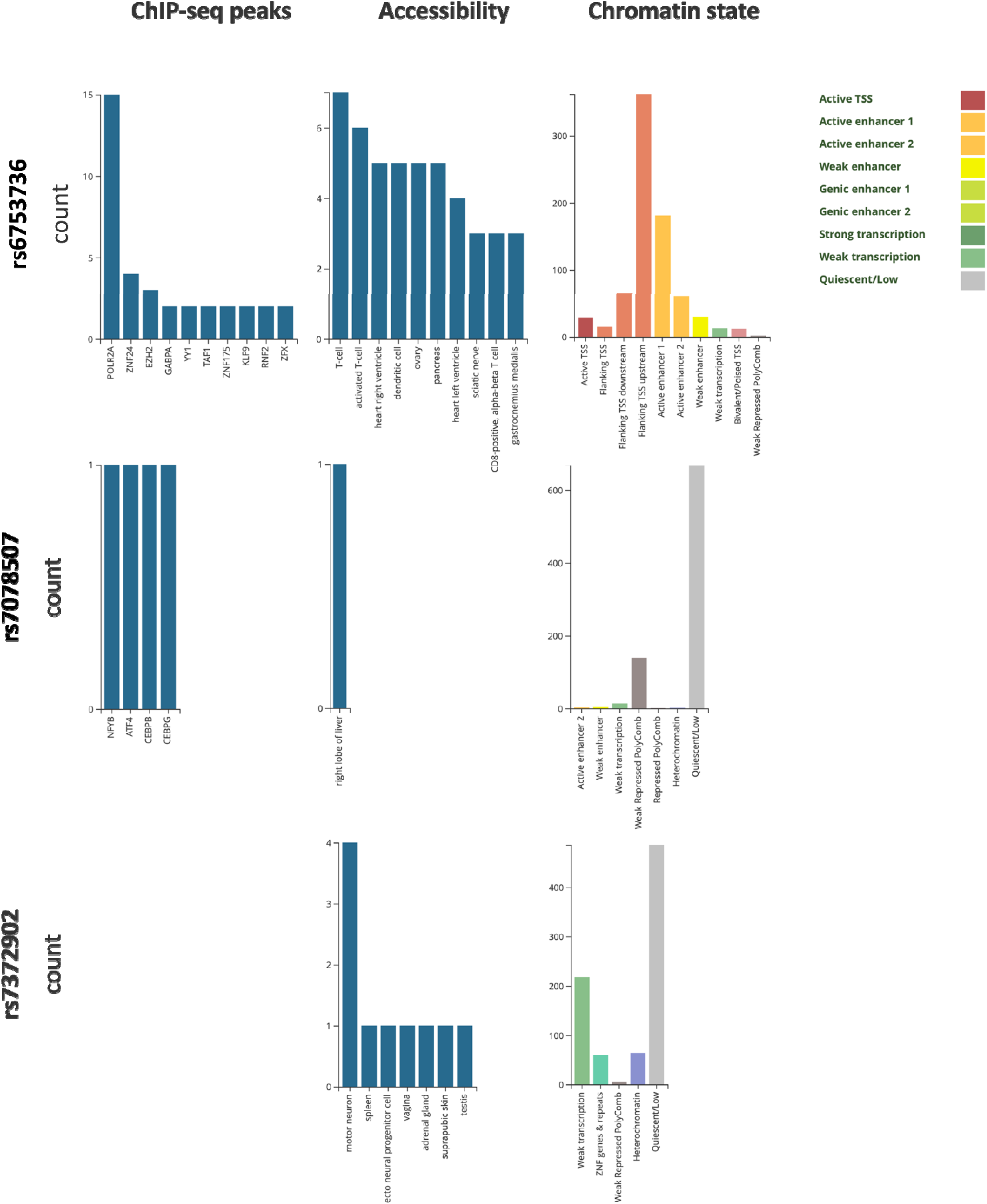
Summary of CVD-associated SNPs from Figure 3 in regulatory regions. Data was downloaded from RegulomeDB [62]

**Table 1:**
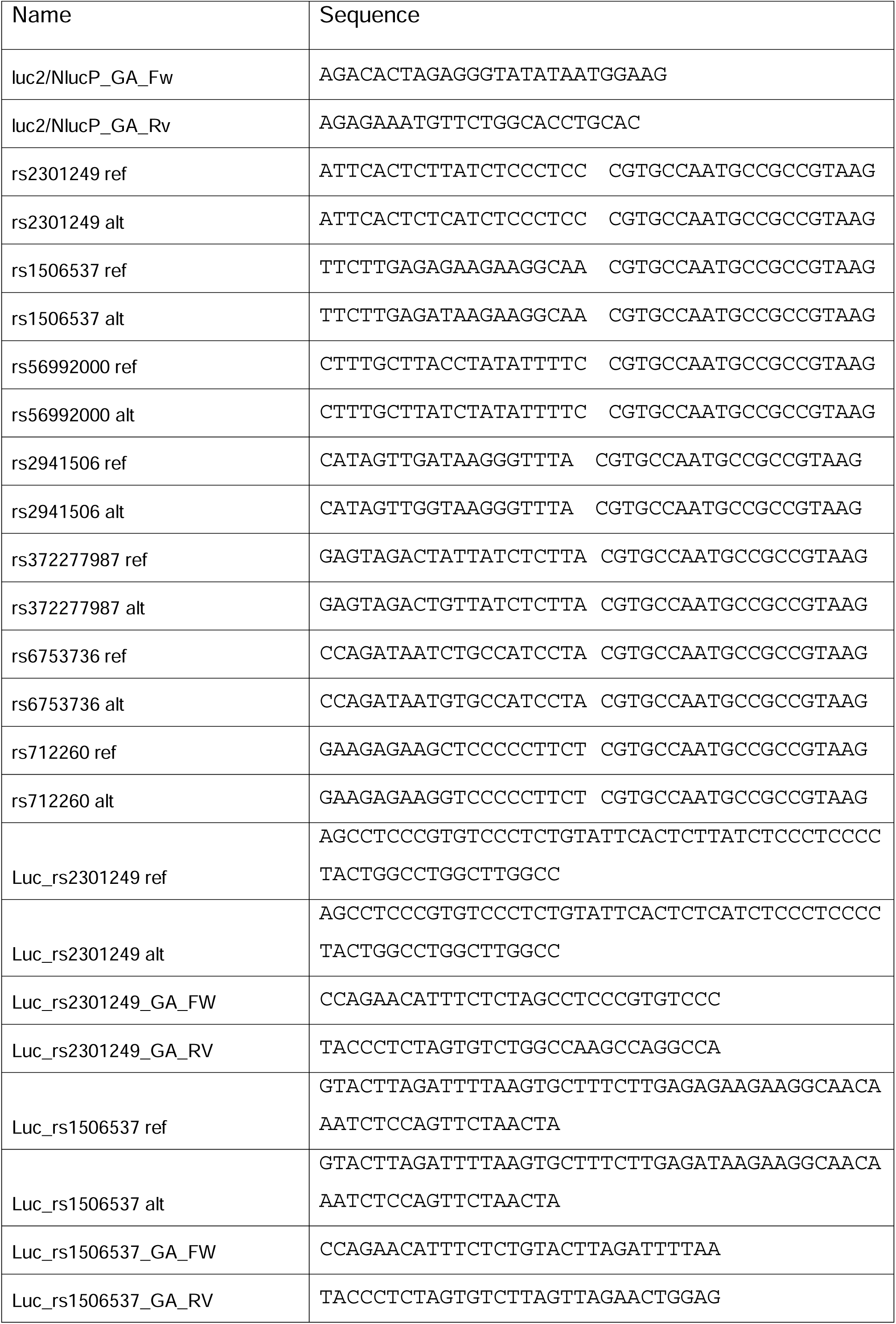
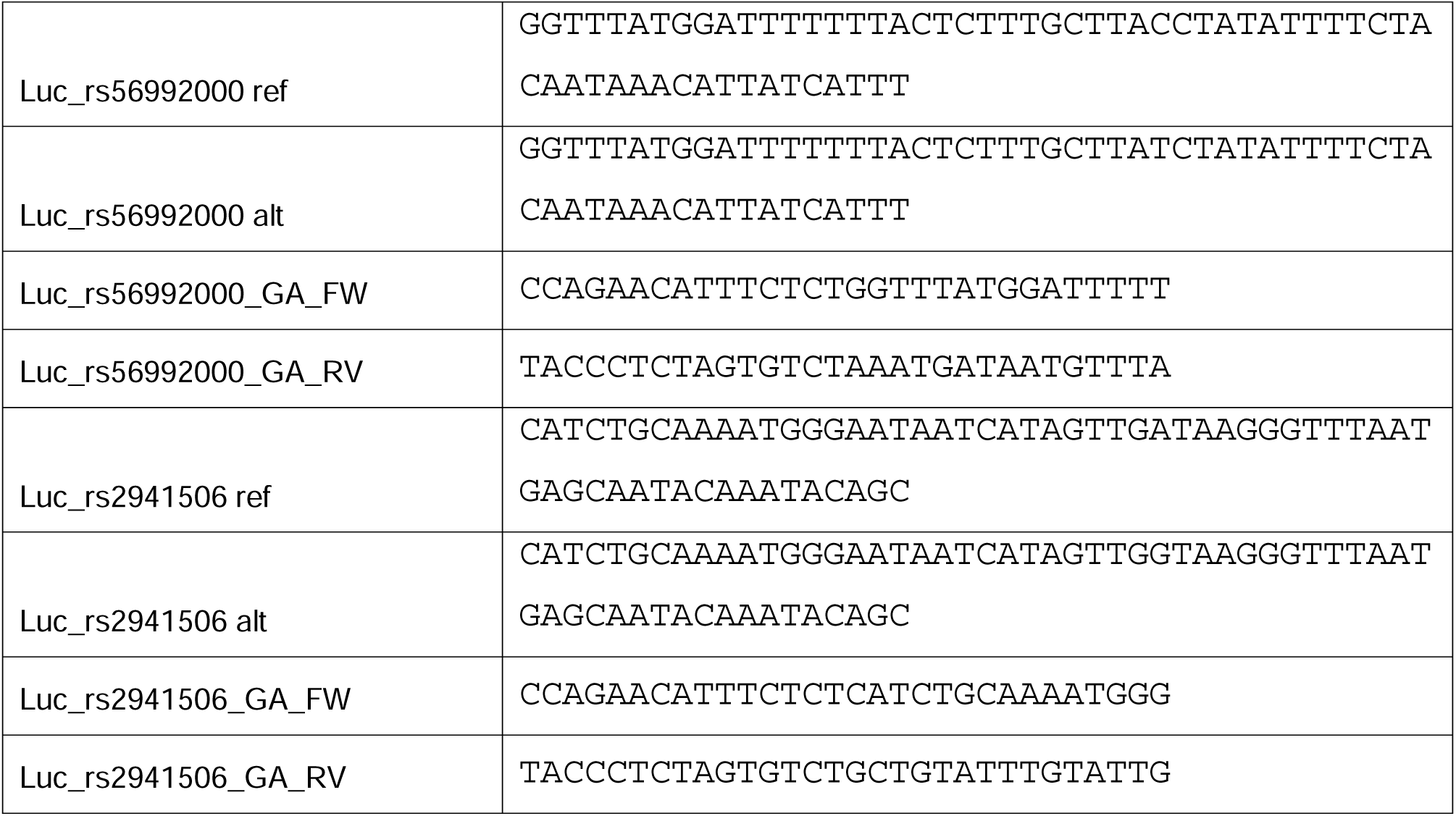
Primers used for cloning and fluorophore labeling active.

## Notes

### Competing Interest Statement

The authors have declared no competing interest.

